# Assembly of the complete horse genome reveals variants associated with reproduction and temperamental traits

**DOI:** 10.1101/2025.09.10.675315

**Authors:** Yu Liu, Xinyu Wang, Hetong Zhang, Ran Yang, Yu Feng, Mo Feng, Tao Yang, Yuanyuan Li, Shiyu Qian, Chunjiang Zhao

## Abstract

A high-quality horse genome assembly is crucial for genetic analysis of equine. Benefiting from long-read sequencing techniques and trio-binning strategy, we assembled a complete telomere-to-telomere horse genome (T2T-horse 1.0) with a size of 2.80 Gb and a base accuracy of more than 99.9999%, comprising a 34.11-Mb entire Y chromosome. T2T-horse 1.0 filled the gaps and corrected errors in previous assemblies, and added 180.30 Mb of previously unresolved regions and 77 new genes compared with the previous reference genome TB-T2T. We identified five types of centromeric satellites and revealed the features of the horse centromeric regions. We characterized the structure of the Y chromosome, and determined the borders of PAR transposed region, and annotated 59 novel genes on Y chromosome of T2T-horse 1.0. Using SNPs in the X-degenerate region of the Y chromosome of T2T-horse 1.0, we performed selection signatures analysis between indigenous horses and horses from intensively selected breeds, and detected the selected genes *ARSF* and *NLGN4Y*, which are closely associated with reproduction and temperamental traits, respectively.

## Introduction

Horse has had deep impact on the human society in many aspects. However, there is a lack of a complete horse genome, contrary to many other mammals^1–5^. In the past, efforts were made to sequence horse genome and some significant progresses have been achieved. In 2009, a 2.5-Gb horse genome with a scaffold N50 of 46.75 Mb was assembled by sequencing the BACs derived from a female Thoroughbred Twilight, and released as a reference genome (EquCab2.0, NCBI accession Number: GCF_000002305.2)^6^. But there are enormous gaps in the genome sequence, and a number of scaffolds failed to assign to the genome. So, an updated genome of Twilight was published in 2018^7^. In the renewed genome, PacBio sequencing techniques were applied, which enabled Scaffold N50 reaching 87.23 Mb, and remarkably improved the continuity of the genome (EquCab3.0, NCBI Accession Number: GCA_002863925.1), though there are still many gaps remained in the assembly. Recently a telomere to telomere (T2T) horse genome derived from a mule was released (TB-T2T, GCA_041296265.1), which were sequenced with ONT, and significantly improved the genomic sequence assembly. However, TB-T2T, like the two previous assemblies, was derived from a female animal, and the Y chromosome is not included. There are still several gaps in TB-T2T, and the complex regions of the genome, such as the sequences of centromeres, subtelomeric regions, and rDNA remain verified.

Several assemblies of horse Y chromosome have been released, but the enormous repeats and complicated sequence structure impeded the complete assembly of the chromosome. A detailed physical map of the horse Y chromosome was developed in an early study^8^. Partial Y chromosomal sequences of Mongolia Horse and Lippizan horse were assembled with genomic data from next generation sequencing, respectively^9,10^. The latest assembly of horse Y chromosome were conducted with BAC and shot-gun sequencing data from a male Thoroughbred Bravo, a half-sib of Twilight, and a 9.5-Mb fragment of eMSY (male specific region of the equine Y chromosome) was generated and contains 52 genes^11^. However, the complete sequence of Y chromosome, especially the heterochromatic regions, pericentromeric regions, and the telomeres are yet to be assembled.

The above assemblies greatly facilitate the genetic studies of the horse by serving as reference genomes, but they left gaps, misassembled regions, and other issues required to be addressed. In the present study, we integrated the reads of ultralong ONT, PacBio HiFi, and Hi-C with a trio-bin strategy, and eventually achieved a telomere-to telomere complete genome assembly of a male Mongolian horse (T2T-horse 1.0) comprising all autosomes and sex chromosomes. We analyzed the sequences in previously unresolved regions (PURs), most of which are in complex fragments of the genome, such as centromeres, telomeres, and the heterochromatic regions of Y chromosome. Using the newly assembled complete horse genome, we conducted population genomics analyses and identified variants in Y chromosome associated with horse reproduction and temperamental traits.

## Results

### Complete genome assembly

A T2T complete genome assembly (T2T-horse 1.0) were generated with ONT reads and HiFi reads, and high-throughput chromatin capture (Hi-C) data was used to scaffold the contigs and anchor them to pseudomolecules (including all the chromosomes except for Y). The analysis of sequence alignment showed that all the gaps left in previous assembly TB-T2T was mainly located in centromeric regions, and they were patched with ultralong ONT reads in the present study. The two previous assemblies derived from female equids do not include Y chromosome (chrY). We assembled the chrY independently using paternal-specific ONT and HiFi reads with a trio-binning design. The detailed comparison of between T2T-horse 1.0 and the two previous horse genome assemblies (EquCab3.0 and TB-T2T) was shown in Table 1.

**Table 1.**
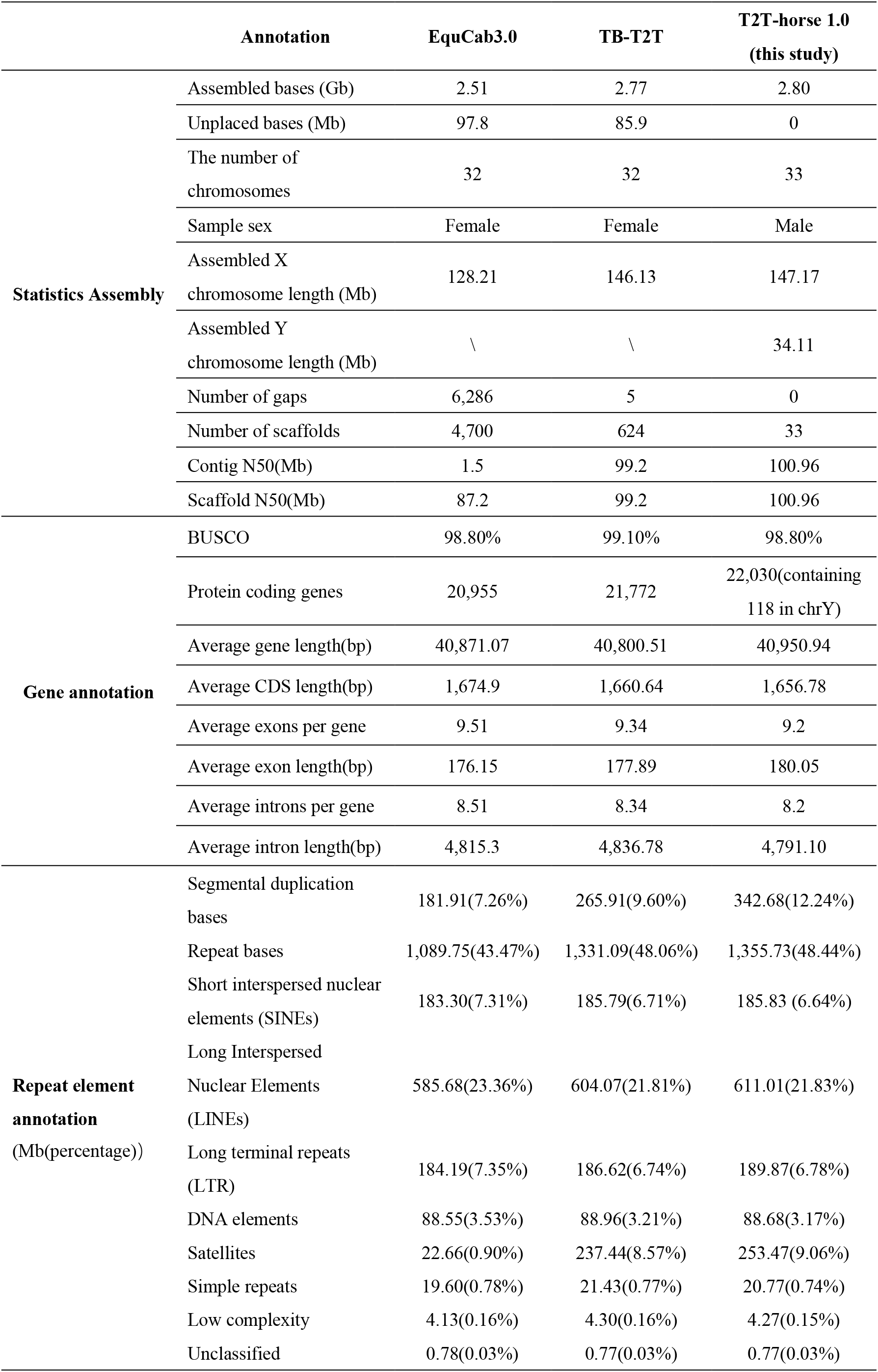
Comparison between EquCab3.0, TB-T2T, and T2T-horse 1.0.

The total size of the complete genome of T2T-horse 1.0 was 2.80 Gb, including a 34.11-Mb Y chromosome. We polished the new assembly and obtained the consensus base quality values 65.28 (>99.9999% base accuracy). There was a high collinearity between TB-T2T (Genbank accession number: GCA_041296265.1) and T2T-horse 1.0. The total size of gaps and misassembled regions (also called previously unresolved regions, PURs) on all autosomal chromosomes and chrX of TB-T2T and EquCab3.0 (Genbank accession number: GCA_002863925.1) are approximately 180.30 Mb and 352.56 Mb (Y chromosome is not included), respectively. The comparison between T2T-horse 1.0 and EquCab 3.0 was shown in Supplementary Fig. 1. The PURs mainly consisted of centromeric satellites and segmental duplications (SDs) (Fig. 1c). T2T horse 1.0 has larger size and more centromeric satellite content than TB-T2T and EquCab3.0. Some assembly errors of TB-T2T were amended in the T2T-horse 1.0. For example, there is a inversion assembly error in chrX in TB-T2T, but they were correctly assembled in the T2T-horse 1.0 proved by alignment of the long reads from ONT and HiFi sequencing (Supplementary Fig. 2).

**Fig. 1.**
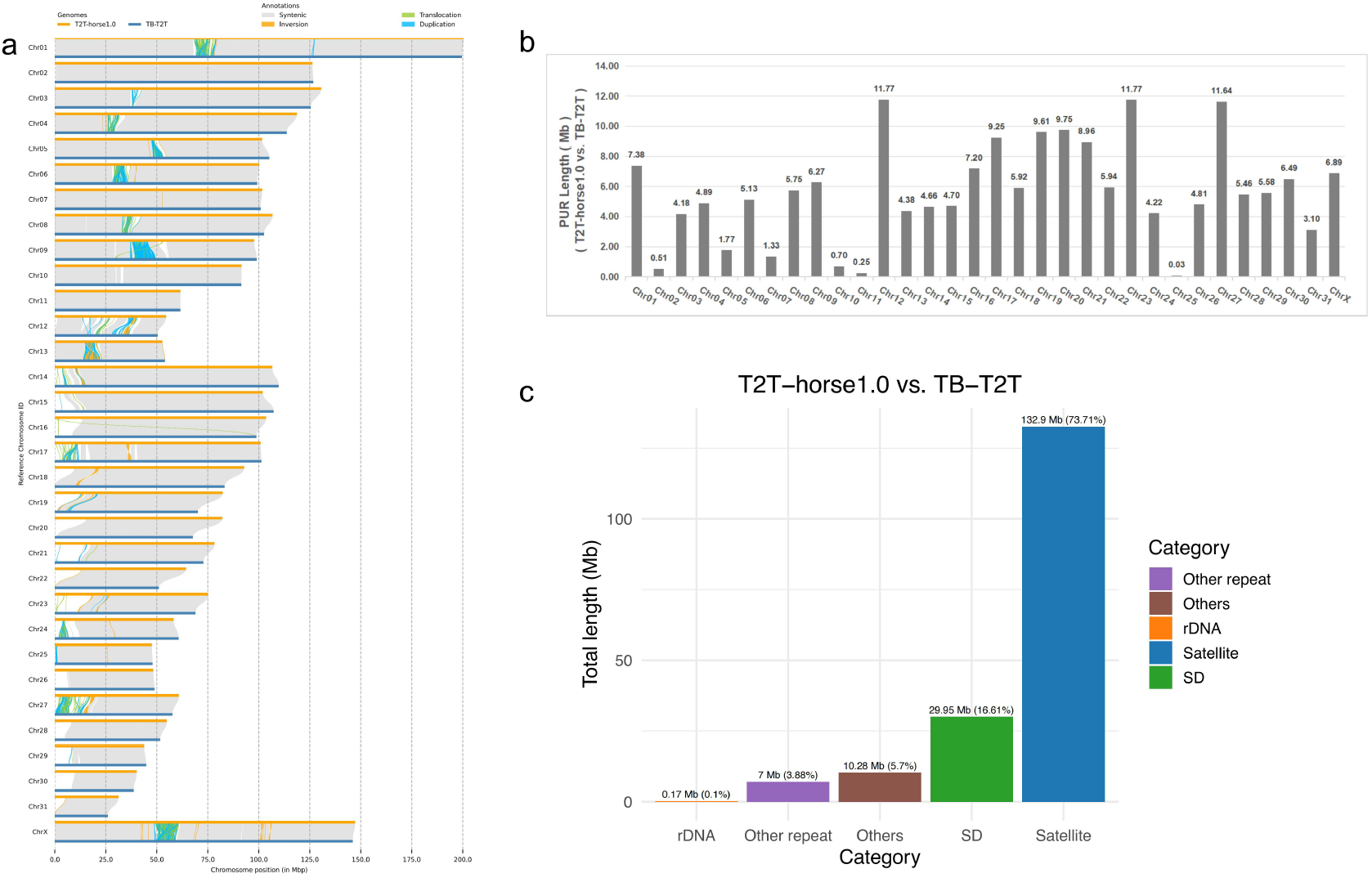
Synteny between T2T-horse 1.0 and TB-T2T and the PURs resolved by T2T-horse 1.0. **a** Synteny between T2T-horse 1.0 and TB-T2T. **b** The PUR sizes in each autosome and chrX of TB-T2T comparing with T2T-horse 1.0. **c** The contents of the PURs in TB-T2T.

### Genome annotation

We identified 22,030 protein coding genes in T2T-horse 1.0, in which 118 genes (containing multiple copies) were located in chrY. For the protein-coding genes in T2T horse 1.0, 315 and 625 genes were identified in the PURs of TB-T2T and EquCab3.0, respectively, and 77 new genes were revealed in the PURs of TB-T2T. Repetitive sequences accounted for 1,355.73 Mb (48.44%) in T2T-horse 1.0, 1,331.09 Mb (48.06%) in TB-T2T, and 1,089.75 Mb (43.47%) in EquCab3.0, respectively. In the repetitive sequences, SDs and LINEs spanned 342.68 Mb and 611.01 Mb, occupying 12.24%% and 21.83% of T2T-horse 1.0, respectively (Table1). In PURs, we added 132.90 Mb satellite sequences and 29.95 Mb segmental duplications (Fig.1c).

### Characterization of Centromeric regions

The centromeric regions were determined with chromatin immunoprecipitation followed by sequencing (ChIP-seq) using anti-centromere protein A (CENP-A) antibody. Centromeres are enriched in satellite repeats, within which CpG sites exhibit high levels of DNA methylation, as confirmed by PacBio HiFi data. Centromeric locations were confirmed with pairwise sequence identity heatmaps. As reported previously^12^, our results from analysis of karyotypes and centromeres showed that there are 13 metacentric or submetacentric chromosomes and 18 acrocentric chromosomes in horse autosomes. Horse chrX is a metacentric chromosome. However, the Y chromosome centromere in horses is acrocentric, whereas it is submetacentric in human, cattle, sheep, cat, dog, and pig^8,13,14^ (Supplementary Fig 3). Centromeric satellite DNA could be grouped into 5 categories: 37cen, 2pi, EC137, sat418, and sat3322 with a repeat motif of 221bp, 132 bp, 137 bp, 418 bp, and 3322 bp, respectively. 37cen is widely distributed in all centromeres except for chromosomes 2 and 11(Fig.2c,d), which is accordant with the previous studies^15^. For example, 37cen constitutes the major component of the centromeric region of chromosome 4(Fig.2a, c). Horse chr11 harbors a satellite-free centromeric region, but we identified 5 copies of 37cen within 2-kb region at the end of chr11 (61,580,647-61,582,643), and they may be the remains of inactivated ancestral centromere. 2pi was reported to be absent from the centromeres of chromosomes 1, 4, 5, 11, 12, and X^15^ and presented in all other Chromosomes. But we found it is only vacant in chromosomes 4, 5, 11 and 12, which was confirmed with FISH (Supplementary Fig 3). According to previous studies^16^, sat418(CENPB-sat) presented in the centromeres of chromosomes 2, 3, 6, 8, 10, 14-25, 27, 29, 30, and 31. Our results overlapped most chromosomes of those from previous studies, but we found sat418 presented in chromosomes X and Y (Supplementary Table4). Sat3322 (satA) was also reported as a centromeric satellite previously^16^, but its distribution remains unknown. In the present study, we found that sat3322 was harbored by chromosomes 2, 26, 31, and the two sex chromosomes (Fig. 2c, d). Our results also indicate that the centromeric region of the Y chromosome is predominantly composed of 37cen, with additional distributions of 2pi, sat418, and sat3322 in the surrounding regions (Fig.2b).

**Fig. 2.**
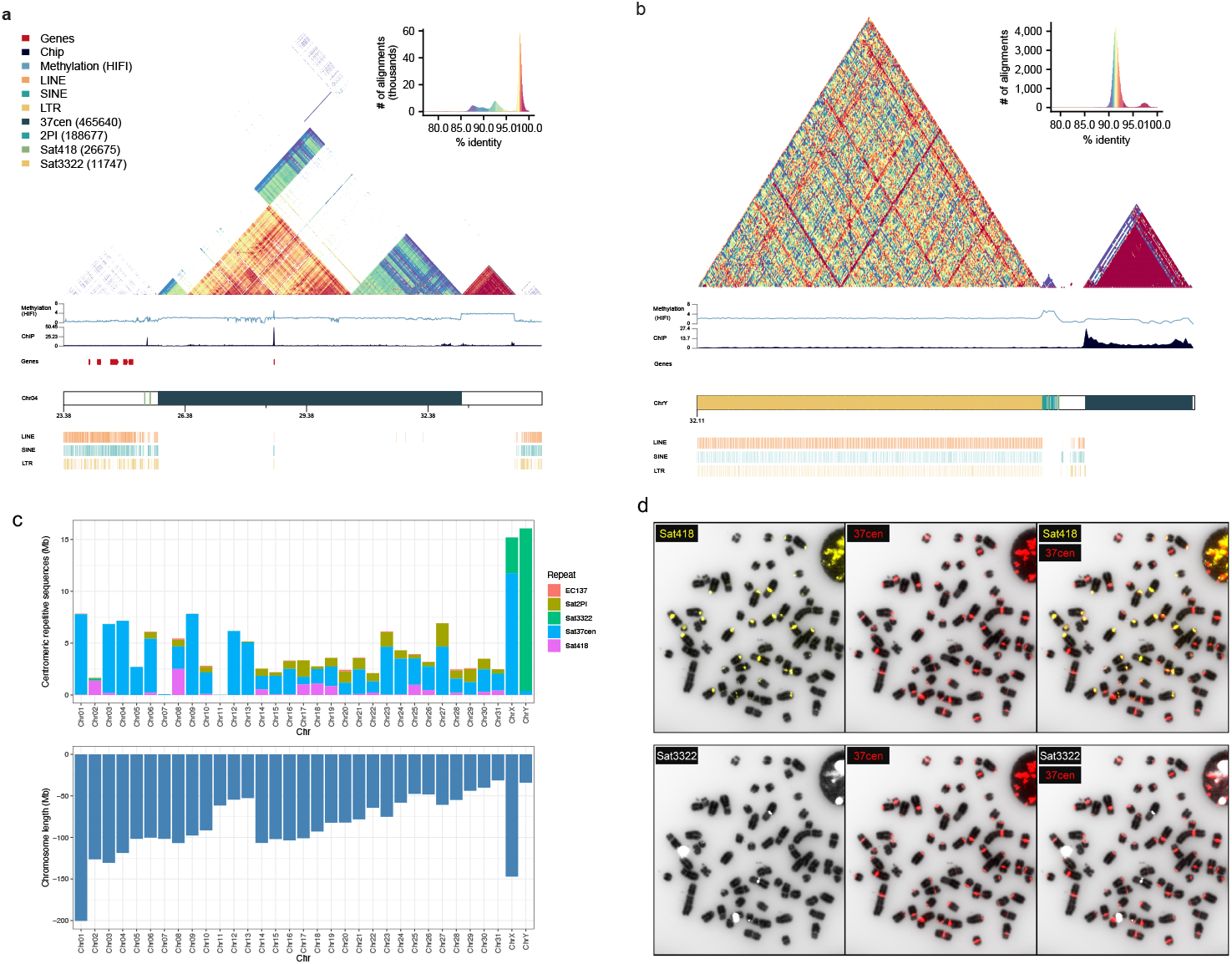
Characterization of centromeric regions. **a** The centromeric region on chr04. The heatmap of sequence identity with color scale is shown across the centromere of chr04. Methylation was determined with HiFi reads. ChIP-seq was performed with the histone H3 variant CENP-A. Genes in the centromere and pericentromeric region are shown in red bars. The centromeric region is occupied by the repeat type 37cen with a few 2PI distributing in the pericentromeric region. Bottom, LINE, SINE, and LTR distributed in the centromere and pericentromeric region. **b** The centromeric region on chr Y. The sequence identity heatmap, ChIP-seq, methylation, and repetitive sequences are shown as described above. No gene was detected in the centromeric region of chrY. Satellites 37cen, Sat3322, Sat418, and 2PI are distributed in the centromere and pericentromeric region. **c** The composition of the centromeric repeat types and lengths of chromosomes. **d** Labelled probes hybridized onto the horse chromosomes.

### Assembly of Y chromosome

The complex sequence structure of chrY and the homology of chrX and chrY hinder the assembly of chrY (Fig. 3a). Compared with the previous horse chrY (eMSY)^11^, T2T horse 1.0 is 24.61 Mb longer (Table 2). T2T horse 1.0 showed good collinearity with the eMSY, but T2T horse 1.0 filled all of the gaps and heterochromatic regions left by the later (Fig. 3b). Compared with the chrY assembly of MH341179, we showed a 5.79% increase in the GC content of T2T-horse 1.0 chrY. Four types of satellites were identified in chrY, comprising 37cen, 2PI, sat418, and sat3322. Our FISH results indicated that sat3322 is mainly harbored in chrY while 37cen constitutes the major content of the centromere (Fig. 2b-d). LINEs, SINEs and LTRs on T2T-horse1.0-chrY have total length of 11.96 Mb, 1.71 Mb, and 3.37 Mb, respectively. Aided with RNA seq expression data, 97 protein-coding genes were annotated on chrY of T2T horse 1.0, in which 59 genes were novel and not annotated previously, and the expression of 5 novel genes (ETY9-13) was validated by qPCR in diverse male and female tissues (heart, liver, spleen, lung, kidney, brain, testis, and ovary) (Supplementary Fig. 4).

**Fig. 3.**
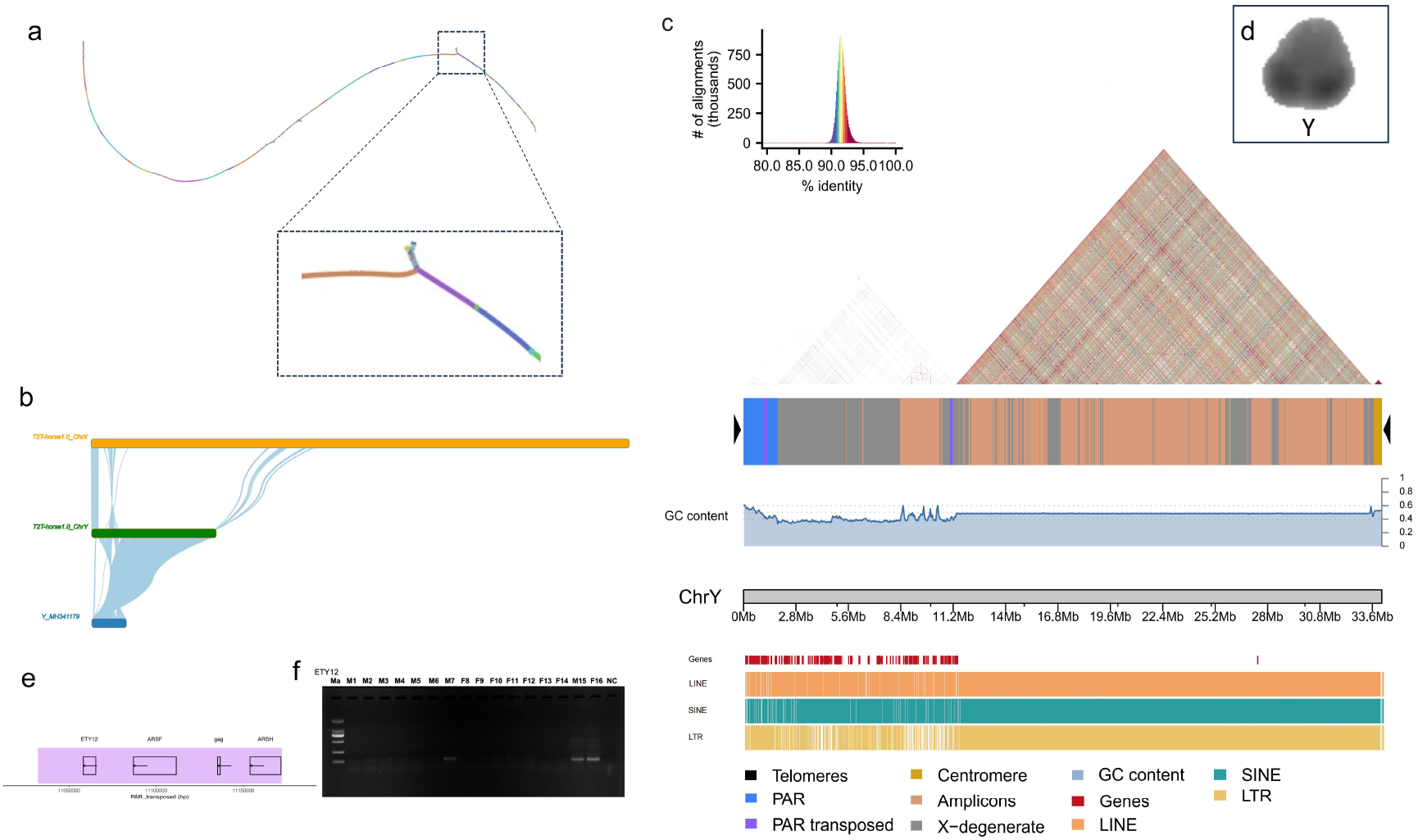
Assembly of horse Y chromosome. **a** A tangle of the assembly between chrY and chrX in PAR. **b** Alignment between T2T-horse 1.0 chrY, T2T-horse 1.0 chrX, and Y_MH341179. **c** The heatmap of sequence identity is shown across chrY. The structure of chrY, GC content and the distribution of genes, LINE, SINE, and LTR are displayed with plot and bars of different colors, respectively. **d** DAPI counterstaining of the Y chromosome. **e** Genes within the par transposed. **f** Reverse transcriptase PCR (RT-PCR) with *ETY12* primers on a panel of 8 adult equine tissues showing testis-specific expression. Ma indicates marker (size standards); M1, male heart; M2, male liver; M3, male spleen; M4, male lung; M5, male kidney; M6, male brain, M7, male testis; F8, female heart; F9, female liver; F10, female spleen; F11, female lung; F12, female kidney; F13, female brain; F14, female ovary; M15, male genomic DNA control; F16, female genomic DNA control; NC, negative control.

**Table 2.**
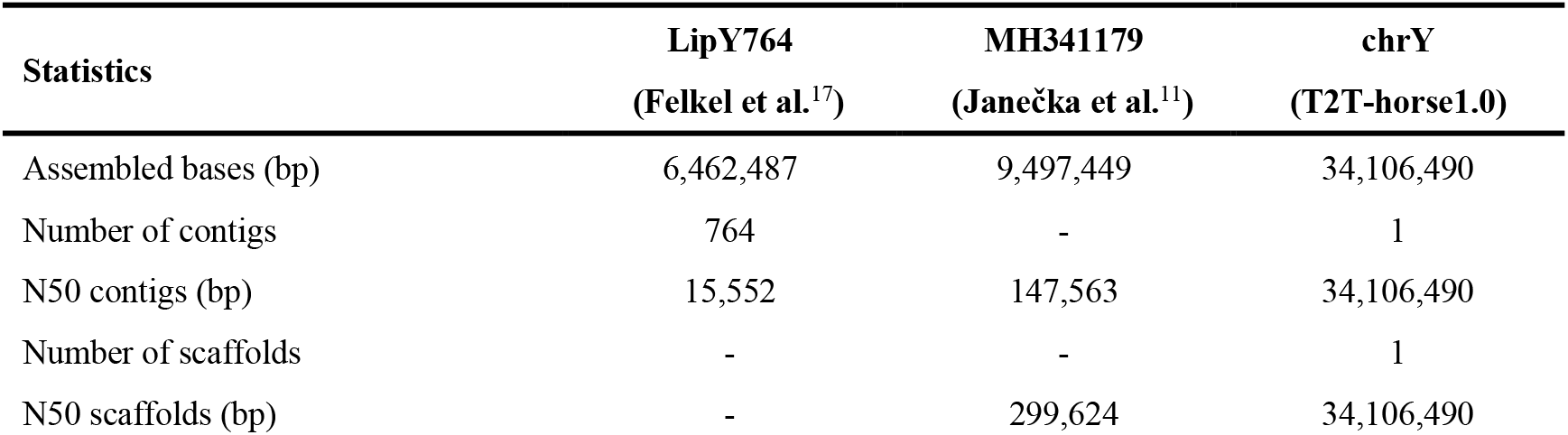
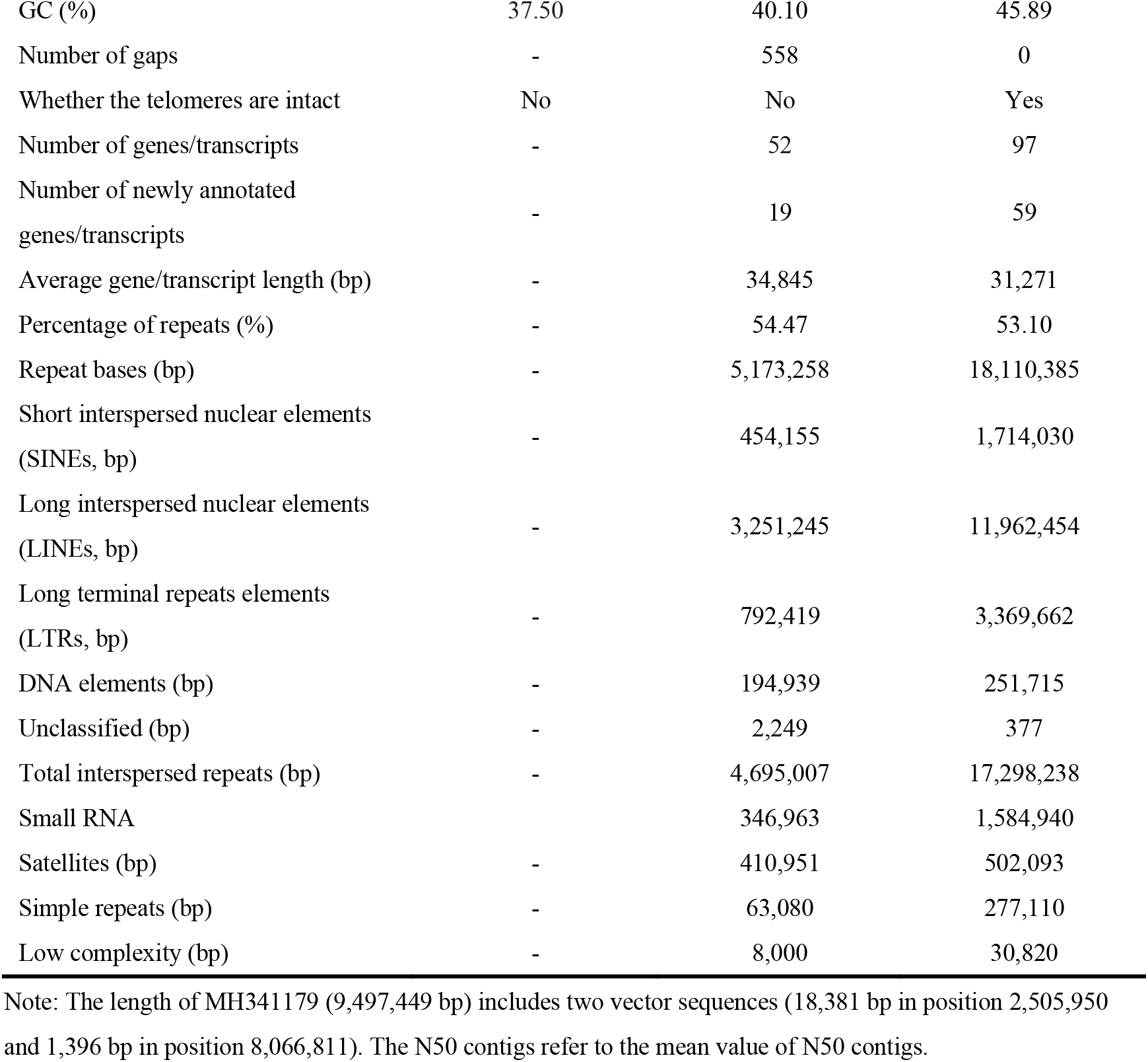
Assembly information of horse Y chromosome.

| Statistics | LipY764<br>(Felkel et al.<sup>17</sup>) | MH341179<br>(Janečka et al.<sup>11</sup>) | chrY<br>(T2T-horse1.0) |
| --- | --- | --- | --- |
| Assembled bases (bp) | 6,462,487 | 9,497,449 | 34,106,490 |
| Number of contigs | 764 | - | 1 |
| N50 contigs (bp) | 15,552 | 147,563 | 34,106,490 |
| Number of scaffolds | - | - | 1 |
| N50 scaffolds (bp) | - | 299,624 | 34,106,490 |
| GC (%) | 37.50 | 40.10 | 45.89 |
| Number of gaps | - | 558 | 0 |
| Whether the telomeres are intact | No | No | Yes |
| Number of genes/transcripts | - | 52 | 97 |
| Number of newly annotated genes/transcripts | - | 19 | 59 |
| Average gene/transcript length (bp) | - | 34,845 | 31,271 |
| Percentage of repeats (%) | - | 54.47 | 53.10 |
| Repeat bases (bp) | - | 5,173,258 | 18,110,385 |
| Short interspersed nuclear elements (SINEs, bp) | - | 454,155 | 1,714,030 |
| Long interspersed nuclear elements (LINEs, bp) | - | 3,251,245 | 11,962,454 |
| Long terminal repeats elements (LTRs, bp) | - | 792,419 | 3,369,662 |
| DNA elements (bp) | - | 194,939 | 251,715 |
| Unclassified (bp) | - | 2,249 | 377 |
| Total interspersed repeats (bp) | - | 4,695,007 | 17,298,238 |
| Small RNA |  | 346,963 | 1,584,940 |
| Satellites (bp) | - | 410,951 | 502,093 |
| Simple repeats (bp) | - | 63,080 | 277,110 |
| Low complexity (bp) | - | 8,000 | 30,820 |

Note: The length of MH341179 (9,497,449 bp) includes two vector sequences (18,381 bp in position 2,505,950 and 1,396 bp in position 8,066,811). The N50 contigs refer to the mean value of N50 contigs.

### The characterization of Y chromosome

We assembled a complete horse Y chromosome with a full length of 34.11 Mb in the present study. Comparing with the T2T-horse 1.0 chrX, we successfully identified the PAR (chrY: 5,032-1,811,313). MSY is located in the region adjacent to PAR and consists of X-degenerate sequences and ampliconic sequences, which were determined with the approach described previously^18^. X-degenerate sequences are mainly located near PAR while ampliconic sequences are distributed next to the X-degenerate, but not neighbor to PAR. We determined the borders of PAR transposed region (chrY:11,032,244-11,171,939), which contains 4 genes: equine testis transcript in Y12 (*ETY12*), arylsulfatase F (*ARSF*), *gag*, and arylsulfatase family member H (*ARSH*). *ETY12* is a newly annotated gene, and only expresses in testis, which was validated using qPCR with several tissues (Fig. 3e, f).

### Ribosomal DNA

We identified rDNA arrays in T2T-horse 1.0, and showed that they are distributed in the subtelomeric region of chromosome 1 and in the pericentromeric regions of chromosomes 28 and 31 in T2T-horse 1.0. We estimated there are 25 rDNA units in haplotype of T2T-horse 1.0 using ddPCR. We detected 17, 5, and 3 copies of rDNA on chr01, chr28 and chr31, respectively, which was validated with FISH. Our results showed that horse has much less rDNA copies than human (N=218, haplotype)^1^ and mouse (N=84, haplotype)^19^. (Supplementary Fig. 5)

### Selection signatures for artificial breeding and reproduction in chr Y

To explore artificial breeding activities, we conducted selection signatures analysis using SNPs in the X-degenerate region of the Y chromosome of T2T-horse 1.0. as the method described previously^20^. We used the SNPs called in the newly assembled chrY to screen the significantly selected genes by comparing 167 indigenous horses (IHs) mainly from East Asian region and 62 horses from intensively selected breeds (SBHs), mostly sport horses. In the region of top 1% windows for F_ST_, there were 2 candidate selected genes, *ARSF* and neuroligin 4 Y-linked (*NLGN4Y*) (Fig. 4a). In the new assembled genome, *ARSF* exists in three similar regions, PAR of chr Y, its homolog in chr X, and a PAR-transposed region in chrY. Remarkably, we confirmed that the top SNP in *ARSF* (chr Y: 11,108,398) was located in the PAR-transposed region by comparing the detailed sequences among the three similar segments (Fig. 4 b, c and Supplementary Fig. 6). The mutation is located in the 7th intron of *ARSF* and there are only 90 bp between the SNP and the 8th exon. We genotyped the loci of 167 IHs and 62 SBHs. Our results showed that the allele G was harbored by 120 IHs while 32 IHs carried allele A, and genotypes of 15 IHs were missing at the locus. The genotyping result from SBHs showed that there are 50 horses carrying allele A and the genotypes of 12 SBHs are missing (Fig. 4d). The SNP of *NLGN4Y* is in intron 1 of the gene (chrY:6,979,863). The genotyping results showed that the allele A, T, and missing were carried by 119, 37, 11 IHs, respectively, while 49 SBHs harbored allele T and the genotypes are missing for the rest 13 SBHs (Fig. 4d). A previous study showed that *ARSF* was associated with reproduction traits in horses^21^. *NLGN4Y* was a gene related with behavioral and cognitive phenotypes in human beings^22,23^.

**Fig. 4.**
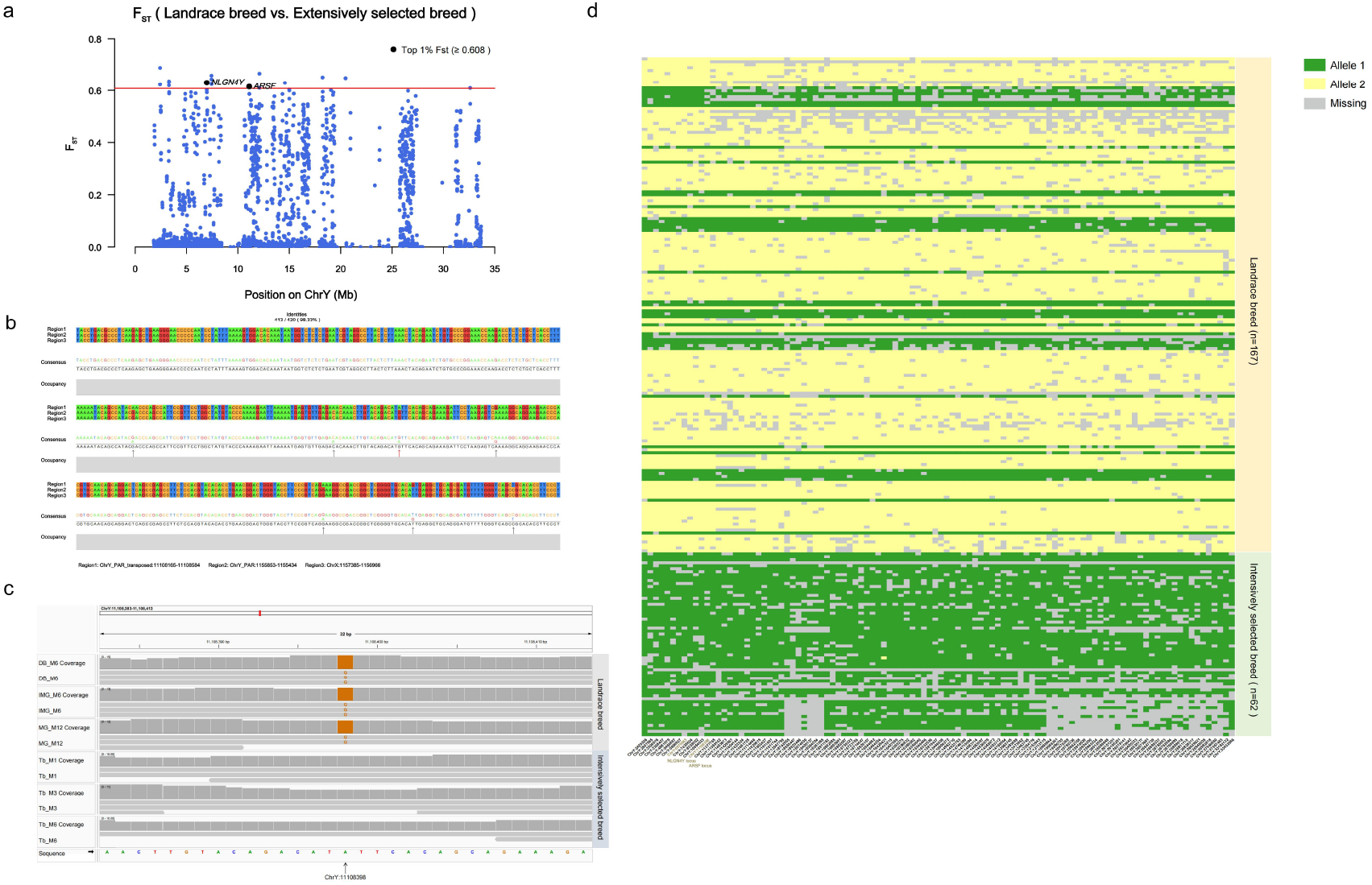
Selection signatures revealed with SNPs in Y chromosome. **a** F_ST_ between indigenous horses and the horses from intensively selected breeds. **b** Alignment of the three homologs of *ARSF*. SNPs are indicated with arrows. **c** The selected SNPs of *ARSF* are shown with the color of orange by comparing three indigenous horses and three horses from selected breeds. **d** Distribution of the haplotype harboring the SNPs of *ARSF* and *NLGN4Y* in indigenous horses and the populations of intensively selected breeds.

## Discussion

The application of the long read sequencing technologies, especially ONT, enables assembly of complete genome^24–26,1,2,27,28,3–5^. In this study, based on the trio-binning strategy successfully applied in previous studies, we assembled a complete T2T-horse genome with entire chrY, and added a new genome in the complete assembly list. In the previous horse assemblies, gaps were mainly distributed in repetitive regions, especially in centromeres and telomeres. We closed the gaps left in previous assemblies with long reads, especially ONT reads, as well as parental data from the trio-binning design, and revealed the complex repetitive sequences in the genome. In the evolution of centromere sequence, satellite repeats are usually absent from centromeric regions of the newly formed chromosomes, but they will be eventually occupied by the repeats to reach mature stage. Unlike donkeys and zebras, more than half of their centromeres vacant of satellite DNA^29,30^, horse chromosome 11 is the only chromosome without satellite sequences at its centromere. Nevertheless, five copies of cen37 were detected at its terminal region, approximately 10,000-fold fewer than those on the X chromosome. This striking difference illustrates the highly variable abundance of the same satellite sequence among chromosomes, and the situation on chromosome 11 is consistent with the model proposed by Eleonora Cappelletti et al.^31^, which suggests that this centromere may be in an immature state with its ancestral centromere gradually being lost. The absence of satellite DNA also facilitated accurate assembly, resulting in strong collinearity and completeness among T2T-horse1.0, TB-T2T, and EquCab3.0 for this chromosome.

ChrY harbors key genes related with male reproduction traits and other genes related to neural development and cognitive traits. After the domestication, artificial selection has posed to domestic horse populations, and modern horse breeds have been subjected to intensive selection, and only a limited number of male horses were selected as stallions based on strict criterions. We compared the intensively selected horses, mainly modern sport horses, with indigenous populations, which did not experience systematic breeding activities, and detected the selection signatures in chrY using the SNPs screened from the newly assembled horse genome. We found that the modern horses harbored a chrY haplotype distinctive from most indigenous horses, which links with some functional genes, for example, the genes *ARSF* and *NLGN4Y*, and they are closely associated with reproduction and temperamental traits^21–23^. The results suggested that these gene loci related with important traits of male horse have been selected in the breeding activities.

In conclusion, T2T horse 1.0 improved the accuracy and continuity of the genome assembly, provides a powerful platform for phylogenetic analysis and study on gene function in horses and related equids.

## Materials and Methods

### Ethical approval

All of the sampling procedures and experimental protocols in the present study were approved by the Institutional Animal Care and Use Committee of the China Agricultural University (AW13014202-1-1).

### Sample collection

The blood samples from a colt at an age of six months and its parents were collected in a Mongolian horse farm in Chifeng city of Inner Mongolia, respectively. The blood samples were stored in −80 °C freezer until the extraction of DNA and RNA. High quality genomic DNA from the samples was extracted. Libraries were constructed as described previously, and the colt was sequenced to obtain 90× next-generation data on the Illumina NovaSeq™ X Plus platform, 74.4× HiFi data on the PacBio Revio platform, and 81.4× (100 k libraries 38.9×, 90 k libraries 42.5×) ultra-long data on the Nanopore PromethION platform. We also sequenced each of the colt’s parents for Illumina 94× and 85.8×. Additionally, 15 tissues collected from Mongolian horse farm in Dezhou city of Shandong were used for RNA-seq (Illumina NovaSeq™ X Plus platform) and Iso-seq (PacBio Revio platform), and then for gene annotation.

### Assembly with trio binning

First, ONT reads were filtered using SeqKit ^32^(v 2.10.1), retaining only those longer than 50 kb. Subsequently, parental short reads were processed with yak (v0.1) (https://github.com/lh3/yak) (https://github.com/lh3/yak) to generate k-mer counts. Assembly was then performed with Hifiasm^33^ (v0.25.0), integrating the filtered ONT reads, the parental short-read k-mers, and HiFi reads, using the following parameters: hifiasm -o horse.asm -t --ul ul.fq.gz -1 pat.yak -2 mat.yak HiFi-reads.fq.gz. Then, we used BLASTN tool^34^ (v.2.10.0) to remove mitochondrial sequences and bacterial contaminants by checking against the NCBI database.

### Construction of Pseudochromosome

Hi-C paired-end reads were processed using HiC-Pro^35^ (v.3.1) (Servant, et al., 2015), which involved aligning them to the draft genome assembly, merging read pairs, and filtering according to restriction fragment sites. The resulting BAM files were then used as input for YaHS^36^ (v1.1), which employed the Hi-C contact information to anchor, order, and orient the contigs, thereby generating a chromosome-scale scaffold assembly. The Hi-C heatmap was visualized using Juicebox^37^ (v.2.18.0) to adjust the contigs of the autosomes and X chromosome.

### Gap filling with ONT reads

Ultralong ONT reads (>50 kb) were first aligned to our assembled genome using Minimap2^38^ (v2.29). The unmapped ONT reads, reads mapped to 1 Mb flanking regions of assembly gaps, and reads with alignment lengths less than half of their original read length were subsequently re-assembled with NextDenovo^39^ (v2.5.2). The resulting contigs were then mapped to the pseudochromosomes using Minimap2^38^ (v2.29), and contigs that uniquely aligned and spanned both sides of a gap were finally used for gaps filling.

### Assembly of chromosome Y

The Y chromosome was assembled using a trio-binning strategy. First, 21-mer and 23-mer databases were constructed from the Illumina short-read sequencing data of the foal and its parents using Jellyfish^40^ (v2.3.0). Parental-specific k-mers were then identified, and paternal k-mers were used to select paternal ONT reads. Reads originating from the autosomes were removed, while those potentially derived from the Y chromosome were retained and corrected with the NextCorrect module in NextDenovo^39^ (v2.5.2). The corrected reads were subsequently used to construct the assembly graph of the Y chromosome with the NextGraph module in NextDenovo^39^ (v2.5.2). The initial Y chromosome assembly was further refined by correcting potential assembly errors using the trio-binning results from Hifiasm. Finally, the Y chromosome sequence was combined with all autosomes and the X chromosome to generate a complete genome assembly.

### Polish of the genome

HiFi reads were aligned to the draft genome assembly using Winnowmap^41^ (v2.03), and the resulting alignments were sorted with SAMtools^42^ (v1.22.1) to generate the HiFi mapping file. For the Illumina short-read data, 21-mer and 31-mer datasets were constructed using Yak (https://github.com/lh3/yak) with k-mer sizes set to 21 and 31, respectively. Finally, the HiFi mapping file together with the k-mer datasets were provided as input to NextPolish2^43^ (v0.2.0) for genome polishing.

### Validation of the complete genome assembly

The assembled genome assembly was validated with the coverage of reads first. Minimap2^38^ (v.2.29) was used to align ONT and HiFi to the new assembly, and the depth coverage in 50 kb windows was estimated with bamdst (https://github.com/shiquan/bamdst) and visualized with R package karyoploteR^44^ (v.1.8.4). Then the T2T assembly was aligned with previous assemblies, TB-T2T and EquCab3.0, using Minimap2^38^ (v.2.29() -ax asm20). Then, SyRI^45^ (v1.7.1) was employed to compare the alignments between the two chromosome-level assemblies in order to identify syntenic regions and structural rearrangements. Visualization of the results was performed using plotsr^46^ (v1.1.0). The quality scores and switch error of the new assembly was estimated with a 21-mer databases derived from the short reads and HiFi reads using Merqury^47^ (v1.3) (Rhie et al, 2020). The completeness of the new assembly was assessed with BUSCO^48^ (v5.8.1) using the laurasiatheria_odb10.

### Annotation for repetitive sequence

RepeatMasker^49^ (v.4.1.1) was applied to annotate repeat with the repeat library derived from the Repbase^50^ of Equus caballus. SDs were annotated using BISER^51^ (v1.4), with filtering criteria applied as described previously^3^.

### Annotation of protein-coding genes

First, EGAPX^52^ (v0.4.0) was used with Illumina RNA-seq data and protein sequences from closely related species (human, pig, mouse, horse, and donkey) as input for initial gene prediction. Subsequently, Iso-seq data were processed with IsoQuant^53^ (v3.7.1) to predict transcript, followed by TransDecoder (v5.7.1) (Haas, BJ. https://github.com/TransDecoder/TransDecoder) to generate gene predictions. In parallel, annotation files from TB-T2T were transferred to our assembly using Liftoff^54^ (v1.6.3). Illumina RNA-seq data were then aligned and assembled, and the resulting short-read transcripts (from StringTie^55^ (v3.0.1)) together with long-read transcripts (from IsoQuant^53^ (v3.7.1)) were integrated with PASA^56^ (v2.5.1) to establish a transcriptome database. These various sources of annotation (EGAPX^52^ (v0.4.0), IsoQuant^53^ (v3.7.1), and Liftoff^54^ (v1.6.3)) were further combined and evaluated using EviAnn^57^ (v2.0.2). PASA^56^ (v2.5.1) was subsequently employed to refine the annotation by adding UTRs and isoforms. Finally, the predicted genes were aligned against the SwissProt protein database using DIAMOND^58^ (v2.0.15) to assign gene names based on conserved protein homology.

### Methylation analysis with long reads

HiFi data were applied to determine 5-methylcytosine methylation across the genome. The HiFi data were aligned to the new assembly with pbmm2(v.1.13.0) (https://github.com/PacificBiosciences/pbmm2). The methylated sites were examined with pb-CpG-tools (v.3.0.0) (https://github.com/PacificBiosciences/pb-CpG-tools).

### Identification of centromeric regions and repeats in centromeres

Centromeric regions were identified and validated through a multi-step procedure. First, candidate centromeric positions were inferred from Hi-C contact heatmaps. Genomic sequences were decomposed into k-mers using KMC^59^ (v3.2.4), and repetitive units were identified with SRF^60^. Second, the assembled repeat units were compared using Minimap2^38^ (v2.29) with the *-ava-pb* option to detect internal redundancies, which were subsequently removed. Third, the non-redundant repeat units were aligned to the whole genome with BLASTN^34^ (v.2.10.0) to determine their genomic distribution. Finally, ChIP-seq data were used to validate the identified centromeric regions.

### FISH

Fluorescence in situ hybridization (FISH) assay was performed according to the method described by Huang et al. (2021) ^61^ with modifications. Probes were amplified using Cy3- or FITC-conjugated primers in a 50 μL reaction system under the following thermal cycling conditions: initial denaturation at 98°C for 3 min; 35 cycles of 95°C for 30 s, 58°C for 30 s, and 72°C for 30 s; a final extension at 72°C for 1 min. The amplified products were purified and dissolved in 10 μL of hybridization buffer (50% formamide, 10% dextran sulfate, 2× SSC). Cells were treated with colchicine (0.1 μg/mL) for 4 h, followed by hypotonic treatment with 0.075 M KCl at 37°C for 10 min, and fixed in a solution of ethanol:glacial acetic acid (3:1) before slide preparation. Prior to hybridization, slides were treated with RNase A at 37°C for 1 h, denatured in 70% formamide/2× SSC at 75°C for 1 min, and then dehydrated through an ice-cold ethanol series. A total of 10 μL of hybridization mixture was denatured at 95°C for 10 min and applied to the denatured chromosomal spreads. Cover slips were sealed with rubber cement, and hybridization was carried out in a humid chamber at 37°C for 16 h. Post hybridization washes included three 5 min rinses in 2× SSC at room temperature and one 10 min wash in 1× PBS. Chromosomes were counterstained with DAPI (0.5 μg/mL) for 5 min and mounted with anti-fade medium. Images were captured using an Olympus BX63 fluorescence microscope equipped with a DP80 camera and processed with Adobe Photoshop CC software.

### Y structural analysis

To identify the pseudoautosomal region (PAR) boundaries, the X and Y chromosomes of the T2T-horse1.0 assembly were aligned using Minimap2^38^ (v2.29) with the parameter -cx asm5. The alignments were visualized with paf2dotplot (https://github.com/moold/paf2dotplot) to facilitate boundary detection. The ampliconic and X-degenerate regions were defined following the approach described in Zhou et al. For the PAR transposed region, the Y chromosome was divided into 5-kb windows, which were aligned using BLASTN^34^ (v.2.10.0) to detect segments sharing more than 95% sequence similarity between the PAR and the PAR-transposed region. The precise boundaries of these regions were then determined by pairwise alignment with nucmer (MUMmer^62^ v4.0.0 software) using the parameters --maxmatch -l 100 -c 500.

### Y-specific SNPs calling and detection of selective sweeps on the Y chromosome

The Short reads of 167 indigenous horses and 62 horses from fine breeds (Supplementary Table 6) were mapped to the Y chromosome of T2T-horse 1.0. All of the samples had an average coverage >14. SNPs of the samples were called with bcftools^42^ (v1.22). Then, we followed the approach described in Rossi C et.al.^20^ to screen SNPs within the X-degenerate region and applied Heng Li’s SNPable (https://lh3lh3.users.sourceforge.net/snpable.shtml) filter for variant filtration, resulting in a total of 10,409 variants retained for downstream analyses. Population differentiation was assessed using the FST method to detect selection signatures between the two populations. The calculations were performed with VCFtools^63^ (v0.1.17). The visualization of results was carried out using the R package qqman (v0.1.9) and Cairo (v1.6-2). Gene frequency was calculated with PLINK1.9^64^, and the visualization of these results was performed with TBtools-II^65^ (v2.337).

## Data availability

The T2T-horse1.0 genome has been uploaded to the NCBI database, with the project number: PRJNA1317574 (http://www.ncbi.nlm.nih.gov/bioproject/1317574).

